# GDF11 secreting cell transplant efficiently ameliorates age-related pulmonary fibrosis

**DOI:** 10.1101/2024.09.06.611670

**Authors:** Li Guo, Pascal Duchesneau, Evan Sawula, Eric D. Jong, Chengjin Li, Thomas K Waddell, Andras Nagy

**Affiliations:** Lunenfeld Tanenbaum Research Institute, Sinai Health System, Toronto, Ontario, Canada; Division of Thoracic Surgery, Toronto General Hospital Research Institute, University Health Network, University of Toronto, Toronto, Ontario, Canada; Institute of Medical Science, University of Toronto, Toronto, Ontario, Canada; Australian Regenerative Medicine Institute, Monash University, Melbourne, Victoria, Australia; Department of Obstetrics & Gynecology, Faculty of Medicine, University of Toronto, Toronto, Ontario, Canada

**Keywords:** Idiopathic pulmonary fibrosis, GDF11, cell and gene therapy, regenerative medicine, anti-aging

## Abstract

Here, we present a combination of cell and gene therapy that harnesses the regenerative properties of GDF11 in age-related pulmonary fibrosis. Our genome-edited FailSafe^TM^-GDF11 mouse ESC line provides controlled proliferation and efficient derivation to lung progenitors while inducibly expressing GDF11. When these cells were transplanted into bleomycin-injured aged mice, they acted as a source of reparative cells, restoring the damaged alveolar epithelium. Furthermore, the transplanted cells acted as an “in situ factory”, enabling the production of GDF11 in response to the inducer drug. This approach attenuated age-associated senescence and led to the successful resolution of fibrosis. Our study presents a promising method for treating pulmonary fibrosis. Additionally, this approach offers a versatile tool that can be expanded to incorporate other regenerative and anti-aging factors. This helps overcome limitations such as high production costs and a short half-life of therapeutic factors. One of the strengths of our system is its ability to allow precise regulation of factor-expression when needed to address specific aging phenotypes.

## Introduction

As we age, our tissues undergo degenerative changes that are closely linked to the development of chronic diseases and cancer. This degeneration becomes increasingly pronounced as our body’s regenerative capabilities diminish, leading to a decline in organ function and overall health. To address this, it is vital to bolster natural regeneration processes that can facilitate injury recovery and slow the advancement of age-related disease conditions.

Discovering, understanding and engineering factors that promote regeneration and suppress degeneration could lead to effective medical treatments for restoring health in situations where natural healing is insufficient. A multitude of factors, including sirtuins^1–6^, Klotho^7–13^, insulin-like growth factor 1 (IGF-1)^14–17^, platelet factor 4 (PF4)^18^ and growth differentiation factor 11 (GDF11) ^19,20,21^ have been proposed as prospective regenerative or anti-aging agents. GDF11, in particular, has been extensively researched for its role in upholding organ function and promoting regeneration in various diseases^19^. However, investigations into GDF11’s efficacy in addressing age-related conditions such as cardiovascular disease, skeletal muscle degeneration, and osteogenesis initially yielded conflicting outcomes ^22–25^.

By now, the controversy surrounding GDF11 has been mostly resolved by discovering tissue-specific variation in its expression and function^19^. Recent reviews have highlighted that the dispute around GDF11 function is presumably rooted in its diverse effects, which depend on the tissue, organ system, and disease-specific disparities in GDF11 expression^26,27^. A further positive development is that there has been a significant increase in publications demonstrating the positive therapeutic effects of GDF11 in treating various conditions^28–32^. The current understanding is that GDF11 is unlikely to be the sole mediator of regeneration, but it does play a role in specific conditions and organ functions.

Idiopathic pulmonary fibrosis (IPF) is a devastating lung disease with limited treatment options. The age-related decline in lung function significantly contributes to the development of IPF^33–35^, reducing the repair capabilities and resolving fibrosis. This leads to irreparable tissue damage, further impaired function, and high mortality rates. Therefore, it is crucial to develop strategies to counteract age-related decline in lung function and promote effective restoration of homeostasis.

The “hallmarks of aging” describe the mechanisms that cause a decrease in regenerative capacity. These hallmarks include stem cell exhaustion, telomere attrition, mitochondrial dysfunction, epigenetic alterations, genomic instability, cellular senescence/apoptosis, defective proteostasis, dysregulated nutrient sensing and distorted intercellular communication^36–38^. Cellular senescence and telomere attrition^39–41^ are two characteristic signs of aging that are especially relevant to humans with IPF. They present promising targets for regenerative and anti-aging interventions. Since it remains to be determined whether GDF11 has regenerative and anti-aging effects on the lungs and age-related pulmonary conditions. Therefore, we have selected GDF11^42,43^ as a prototype for our study targeting IPF.

Like other protein-based regenerative molecules, the practical usage of *in vitro* manufactured recombinant GDF11 (rGDF11) is hampered by its high cost and short half-life (12h)^44^. Moreover, like many other hormones, GDF11’s beneficial effects are only seen in a narrow concentration window. Excessive levels of GDF11 could potentially lead to adverse consequences, such as neurotoxicity, cachexia and mortality^45–47^. These limitations highlight the need to explore alternative approaches.

An alternative method involves using disease-site-positioned therapeutic cell grafts engineered to secrete GDF11 in response to small-molecule inducers. This approach could specifically target a particular organ or site. The biological activity of GDF11 is dependent on various factors such as disease pathologies, organ function, affected tissue and the aging process^19,26,48^. Therefore, a cell transplant-based system that allows for controlled expression of GDF11 is feasible and necessary to achieve therapeutic benefits while minimizing systemic adverse effects. It also permits the dose and duration optimization of the treatment.

We have established ESCs expressing GDF11 in a doxycycline-inducible manner. The cells also contained our FailSafe^TM^ (FS) system that ensures safety by eliminating harmful, highly proliferative cells through ganciclovir (GCV) treatment prior to or post transplanting lung progenitors differentiated *in vitro* from these genome-edited ESCs^49^.

We used the bleomycin (BLM) induced injury model, which recapitulated some of the critical features of the age-associated lung deterioration of IPF patients. We demonstrated that these cells could integrate to replenish the pool of alveolar progenitors to resolve the disrupted alveolar epithelium and functionally contribute to tissue regeneration. More importantly, the engrafted cells are an in situ “factory” of GDF11, efficiently attenuating age-associated senescence for successful fibrosis resolution.

This study provides a new therapeutic approach to treating age-related pulmonary fibrosis by combining cell and gene therapy. The approach also serves as a therapeutic model for other degenerative diseases, where the use of *in vitro*-produced clinical grade biologics is limited by high cost, short half-life and frequent treatment needs.

## Results

### 1. GDF11 expression declines in aging lungs under both physiological health and pathological conditions

In both adult humans and mice, the expression of GDF11 varies across different tissues at both the mRNA and protein levels^19^. Therefore, we first evaluated GDF11 expression in the lung during aging under physiological health and pathological conditions.

In young (8-10-week) and two old mice groups (12-month and 24-month), Gdf11 lung expression levels were compared using qRT-PCR. While the young mice’s expression level was significantly higher than the old, the two old groups did not differ significantly. Therefore, the 12-month age was chosen for the “old” group (Fig. 1a) in the studies presented. The qRT-PCR analysis-based findings were supported by immunohistochemistry, showing the decreased level of GDF11 protein in the distal lung (Fig. 1b,c).

**Figure 1.**
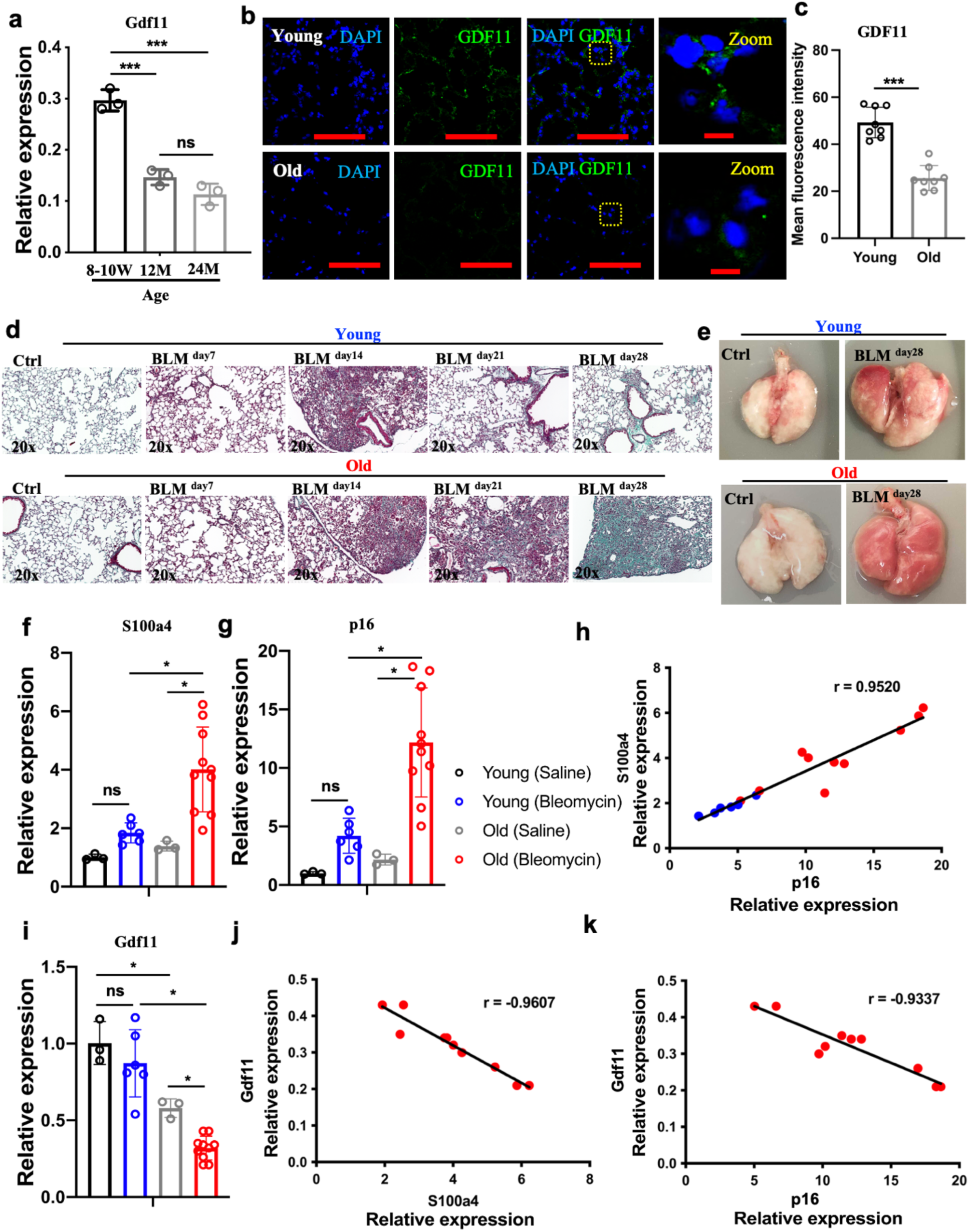
GDF11 expression declines in aging lungs under both physiological health and pathological conditions **a** The expression of GDF11 in the lungs from the mice of different ages, as measured by qRT-PCR; **b** Confocal microscopy images of distal airways of young mice (8-10 weeks) and old mice (12 months) showing nuclear stain DAPI (blue) and GDF11 (green); **c** Quantification of mean fluorescence intensity of GDF11; **d** Representative images of Masson’s trichrome staining of the lungs of young and old mice after 7, 14, 21, and 28 days of BLM-induced pulmonary fibrosis. Interstitial collagen (blue) still persisted in the lungs of old mice 28 days after BLM treatment; **e** Representative images of whole lungs from all the experimental groups at days 28 after BLM treatment; The expression of the fibrosis gene S100a4 **f** and the senescence marker p16 **g** in the lungs of young and old mice 28 days after saline or BLM treatment, as measured by qRT-PCR comparing fold differences in gene expression in young, healthy controls; **h** The correlation of the expression of S100a4 with p16; **i** The expression of GDF11 in the lungs of young and old mice 28 days after saline or BLM treatment, as measured by qRT-PCR comparing fold differences in gene expression in young, healthy controls. The correlation of the expression of GDF11 with S100a4 **j** and p16 in the group of old mice **k**. *p < 0.05; **p < 0.001; ***p < 0.0001. Scale bar, 100 μm (**b**);10 μm (**b**-zoom).

BLM-induced injury predominantly targets the distal lung, leading to basement membrane disruption and fibrosis development in the alveolar compartment^50^. While not an accurate model of human IPF, this model mimics age-related deteriorations in lung function, including enhanced p16 activation indicative of senescence^51^. Our study corroborated earlier findings^52^ by demonstrating that old mice exhibited impaired fibrosis resolution compared to the nearly complete recovery seen in young mice on day 28 post-BLM injury (Fig. 1d, e, f). Thus, given the lingering fibrosis in old mice, we selected day 28 to evaluate lung GDF11 expression.

Age-dependent accumulation of senescence has been implicated in human IPF^51^. Our study revealed sustained elevation of the senescence marker p16 in old mice at day 28 post-BLM administration compared to young mice (Fig. 1g). Additionally, we observed a strong correlation (r = 0.9520) between p16 levels and the fibrosis marker S100a4 expression (Fig. 1h), indicating a close relationship between senescence and fibrosis severity.

Compared to healthy controls, BLM-injured old mice showed decreased GDF11 levels, suggesting an accelerated age-related decline of Gdf11 due to BLM injury (Fig. 1i). Additionally, we found significant negative correlations between Gdf11 levels and S100a4 expression (r = -0.9607) (Fig. 1j) and p16 (r = -0.9337) (Fig. 1k). These findings indicate that GDF11 levels decrease in aging lungs under both normal physiological and BLM-induced pathological conditions. This reduction is associated with the impaired fibrosis resolution mediated by cellular senescence.

### 2. Exogenous GDF11 protein administration partially ameliorates age-related deterioration of fibrosis resolution in the distal lung

Alveolar type II cells (AEC-IIs) in the distal lung, akin to stem cells in other tissues^53,54^, show diminished regenerative capability with age. This decline, exacerbated by protein homeostasis disruptions, results in cellular harm and tissue dysfunction^55,56^. Our investigation into aging’s impact on AEC-IIs began with analysing surfactant protein C (SPC) expression, a crucial protein for AEC-IIs maintenance. As anticipated, old lungs exhibited reduced SPC protein levels (Fig. 2a).

**Figure 2.**
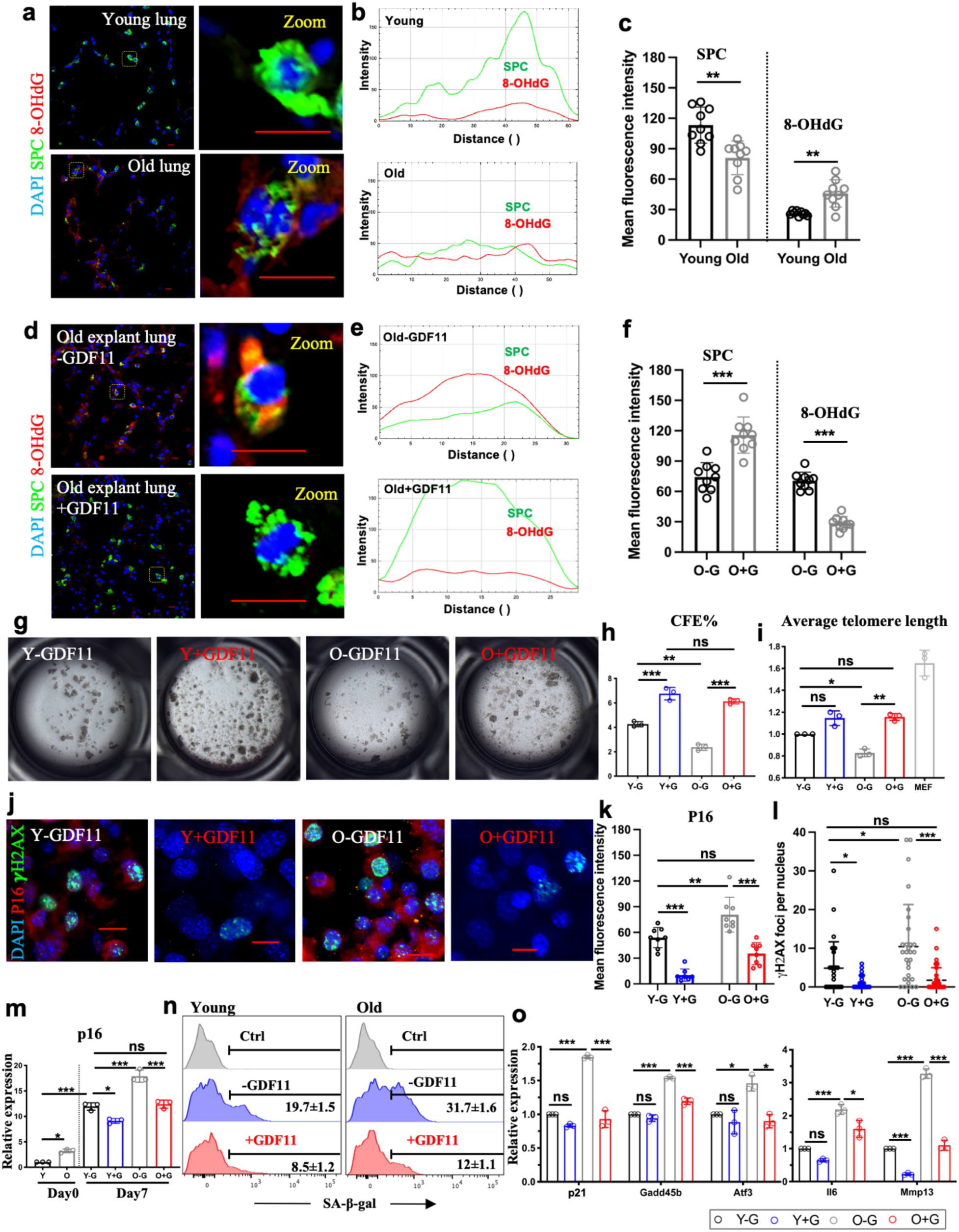
Exogenous GDF11 can partially ameliorate age-related cellular deterioration in the distal lungs **a** Confocal microscopy images of distal airways of young mice (8-10 weeks) and old mice (12 months) showing nuclear stain DAPI (blue), SPC (green) and 8-OHdG (red); **b** Fluorescence intensity profile plot of zoomed cells; **c** Quantification of fluorescence intensity of SPC and 8-OHdG; **d** Representative confocal microscopy images of distal airways of old explant lungs treated with or without GDF11 protein showing nuclear stain DAPI (blue), SPC (green) and 8-OHdG (red); **e** Fluorescence intensity profile plot of zoomed cells; **f** Quantification of fluorescence intensity of SPC and 8-OHdG; **g** Bright-field images depicting the generation of colonies derived from AEC-IIs isolated from young and old mice in the absence and presence of GDF11 recombinant protein; **h** The colony-forming efficiency; **i** Expression levels of average telomere length in cells obtained from each condition, as measured by qRT-PCR comparing fold differences in the expression in control young cells; **j** Representative confocal microscopy images of AEC-IIs isolated from young and old mice in the absence and presence of GDF11 recombinant protein showing nuclear stain DAPI (blue), ***γ***H2AX (green) and p16 (red); Quantification of fluorescence intensity of p16 (**k**) and ***γ***H2AX foci (**l**); **m** Expression levels of p16 in the AEC-IIs obtained from each condition, as measured by qRT-PCR comparing fold differences in the expression in day 0 freshly AEC-II cells of young mice; **n** Flow cytometric analysis of SA-β-gal expression levels in cells cultured in each condition; **o** Expression levels of age-related stress response genes in the p53 tumor suppressor pathway in cells obtained from each condition, as measured by qRT-PCR comparing fold differences in the expression in control young cells. *p < 0.05; **p < 0.001; ***p < 0.0001. In **n**, data are representative of a minimum of three independent biological replicates. Scale bar, 10 μm (**a**, **d** and **j**).

Moreover, we explored age-related mitochondrial DNA (mtDNA) damage, known to impact the reduced lifespan of lung progenitor cells^57–59^. The accumulation of mtDNA damage is a hallmark of aging and is linked to various age-related pulmonary diseases, such as IPF, chronic obstructive pulmonary disease (COPD), and lung cancer. The mtDNA damage induces cellular senescence and apoptosis, contributing to dysfunction in AEC-IIs and the progression of these diseases^58,59^. To assess the extent of mtDNA damage in the aging distal lung, we conducted an immunohistochemical analysis using an 8-hydroxy-guanosine (8-OHdG) antibody to label DNA damage in both the mitochondrial and nucleus^60^. The results showed a significant increase in the number and immunoactivity of 8-OHdG+ cells in the old lungs, indicating that mtDNA damage accumulates with aging in the distal lung, including the AEC-II compartment (Fig. 2a-c).

We found that treatment with recombinant GDF11 halted the age-related decrease in SPC expression (Fig. 2d-f), with no detectable abnormalities (Fig. S2c). Regarding the mtDNA damage, we observed that in the explanted cultured lungs, 8-OHdG expression increased compared to fresh lungs, indicating a worsening of mtDNA damage due to the oxidative stress of the culture environment. Then, we treated the old-lung explants with 50ng/mL^21^ of GDF11 recombinant protein daily for up to 7 days (Fig. S2a). Although treatment with GDF11 for 3 days did not show significant effects (Fig. S2b), remarkably, after 7 days of GDF11 treatment, immunoactivity of 8-OHdG was greatly reduced in the distal lung and SPC+AEC-II cells, suggesting GDF11’s potential in alleviating mtDNA damage from aging and culture-induced oxidative stress.

As many hallmarks of aging in chronic lung diseases are linked to AEC-II cells, reversing their aging traits could potentially retain the regenerative capability of the young in the aging lungs. Therefore, we evaluated the self-renewal potential of AEC-II cells obtained from young (8-12-week-old) and old (12-month-old) mice using a colony-forming assay^61^. AEC-II cells from old mice showed decreased clonogenicity, but treatment with GDF11 protein significantly improved colony-forming efficiency in both age groups (Fig. 2g, h).

Given that telomere shortening contributes to age-related decline in self-renewal and cellular senescence^62,63^, we investigated whether the telomeres were shortened in old AEC-II cells. Indeed, our analysis showed that both the distal lung tissues (Fig. S2d) and AEC-IIs (Fig. 2i) from old mice had shorter telomeres than those from young. Treatment with GDF11 led to significant telomere elongation in aged AEC-II cells. In contrast, there was no significant change in telomere length in young AEC-II cells treated with GDF11, suggesting that the improved clonogenicity of the young may result from reduced *in vitro* oxidative stress (Fig. 2i).

Genomic integrity is a crucial factor in cellular health, especially in the presence of aging-related genetic abnormalities^64^. Repairing DNA double-strand breaks during DNA replication or caused by environmental factors is a significant challenge. Our immunofluorescence analysis of phosphorylated histone H2AX (γH2AX) in lung cells, as a marker of double-strand breaks that increase with aging^64^, revealed a significant rise in both the number and intensity of positive foci in aged lung cells. Interestingly, the treatment with GDF11 reduced the level of γH2AX immunoreactivity in these cells compared to untreated cells (Fig. 2j, i).

Furthermore, GDF11 successfully reduced p16 expression and the senescence marker SA-β-gal activity during *in-vitro* culture of lung tissue (Fig. 2k, m, n). Treatment with GDF11 had a similar effect on stress response genes such as p21, Gadd45b, Atf3, Interleukin-6 (IL6), and metalloprotease Mmp13 in the p53 tumour suppressor pathway. We detected an increase in the expression of these genes in the old lung that was partially reversed by GDF11 treatment (Fig. 2o).

Clearing senescent cells through apoptosis induction has been shown to aid in resolving fibrosis^41^. Therefore, we investigated whether GDF11’s ability to counteract senescence in lung cells is linked to inducing apoptosis in senescent cells or inhibiting cellular senescence. Flow-cytometric analysis revealed a higher percentage of Annexin^+^ PI^-^ cells in cultured old lung cells (36.0%±3.4%) than in young cells (9.2%±0.3%), indicating increased susceptibility to apoptosis with age, likely due to oxidative stress from culture conditions. Upon treatment with GDF11, both young and old cells showed a decrease in apoptosis (young group by 5.5%±0.8%; old group by 18.6%±1.2%) (Fig. S2e), suggesting that GDF11 inhibits senescence instead of inducing apoptosis of the senesced cells.

Taken together, our pilot study using GDF11 recombinant protein showed that GDF11 mitigated some of the age-related deteriorations in the lung

### 3. Generation of lung progenitors with regulatable expression of GDF11

Our FailSafe ESC (FS) line^49^ was engineered with doxycycline (dox) inducible GDF11 transgene (Fig. 3a, b) using piggyBac transposon technology^65^. After screening 14 clonal lines for dox inducibility and differentiation efficiency into definitive endoderm and lung progenitors, the most effective clone was chosen for further studies and designated as FS-GDF11 cells. The FS system ensures safety by eliminating unwanted dividing cells pre- and post-transplantation, depending on the timing of the GCV treatment^49^.

**Figure 3.**
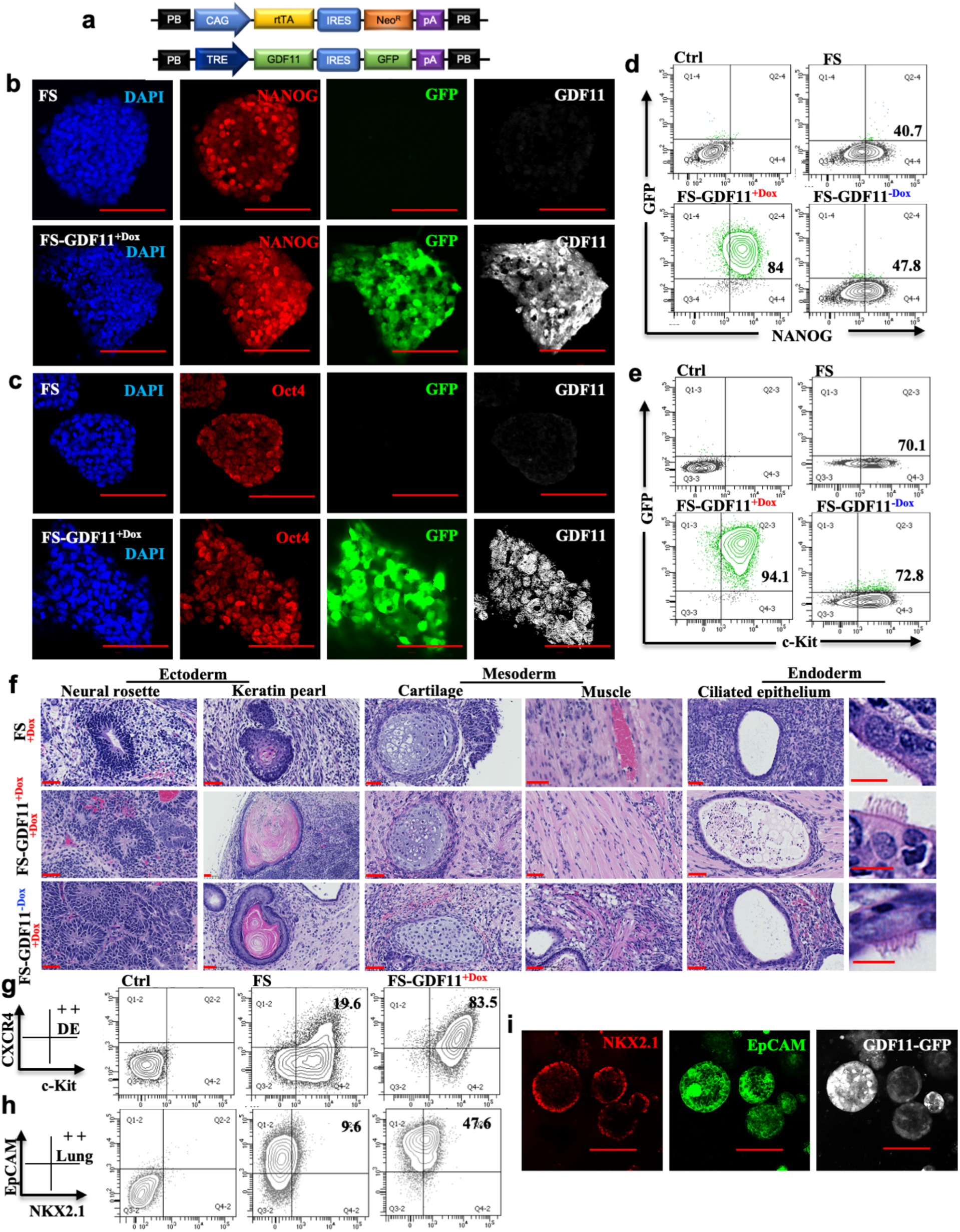
Generation of lung progenitor cells with regulatable production of GDF11 in *vitro*. **a,** FailSafe^TM^ mouse ESCs were genetically engineered with a piggyBac transposon to overexpress GDF11 with a GFP fluorescent reporter. Confocal microscopy images of parental FS and FS-GDF11^+dox^ ESCs, showing cells stained for NANOG **b** and Oct4 **c** pluripotency markers, GFP reporter and GDF11. Representative flow cytometry contour plots of NANOG **d** and c-kit **e** expression levels in parental FS and FS-GDF11 ESCs cultured in the present or absent of dox. **f,** Hematoxylin and eosin staining showed the tissue composition of the teratomas, including ectoderm (neural rosette, keratin pearl), mesoderm (cartilage, muscle) and endoderm (ciliated epithelium). Representative flow cytometry contour plots of c-Kit+CXCR4+ cells representing definitive endoderm **g**, NKX2.1+EpCAM+ lung progenitors **h,** during the differentiation of parental FS and FS-GDF11 mouse ESCs. **I,** Confocal microscopy images showing lung progenitor cells derived from FS-GDF11 ESCs stained with the nuclear stain NKX2.1(red), EpCAM (green) and GFP reporter (grey). In **d**, **e**, **g** and **h**, data are representative of a minimum of three independent biological replicates. Scale bar, 100 μm (**b**, **c** and **i**); 50 μm (**f**).

We analyzed the pluripotency of FS-GDF11 ES cells with and without dox-induced activation of exogenous GDF11 in both in *vitro* and in *vivo*. In *vitro,* all ES cell lines expressed pluripotency-related markers, such as Nanog (Fig. 3b, d), Oct4 (Fig. 3c) and c-Kit (Fig. 3e). In *vivo*, the cells formed teratoma in all mice utilized, displaying all three embryonic germ layers. (Fig. 3f). More importantly, these cells efficiently differentiated into CXCR4+, c-Kit+ definitive endoderm and Nkx2.1+, EpCAM+ lung progenitors (Fig. 3g-i).

### 4. Exogenous GDF11 alleviates age-related and bleomycin-induced aggravated cellular senescence

Hence, we conducted an *in vitro* experiment utilizing BLM-induced accelerated cellular senescence^39,41^ to study the potential anti-senescence properties of different sources of GDF11 on aging lung cells. Of the sources, there were recombinant proteins (rGDF11) and two cell types, such as mouse embryonic fibroblasts (MEFs), which naturally express a certain level of endogenous GDF11^66^ and the FS-GDF11 cells.

CD45-CD31-cells were freshly isolated from the distal region of the lung, which are the main targets of BLM in alveolar epithelial and fibroblast cells. The cells were then seeded on top of the transwell membrane and allowed to recover for two days before subsequent procedures (Fig. 4a). To simulate BLM-injury pathological condition, cells were exposed to a sublethal concentration of BLM (50 ug/mL) on day 3 for 2 days. Then, the cells were maintained in culture for an additional 3 days, either in the presence or absence of rGDF11 or GDF11-producing cells. Concurrently, we examined the response of non-injured cells to these treatments (Fig. 4b).

**Figure 4.**
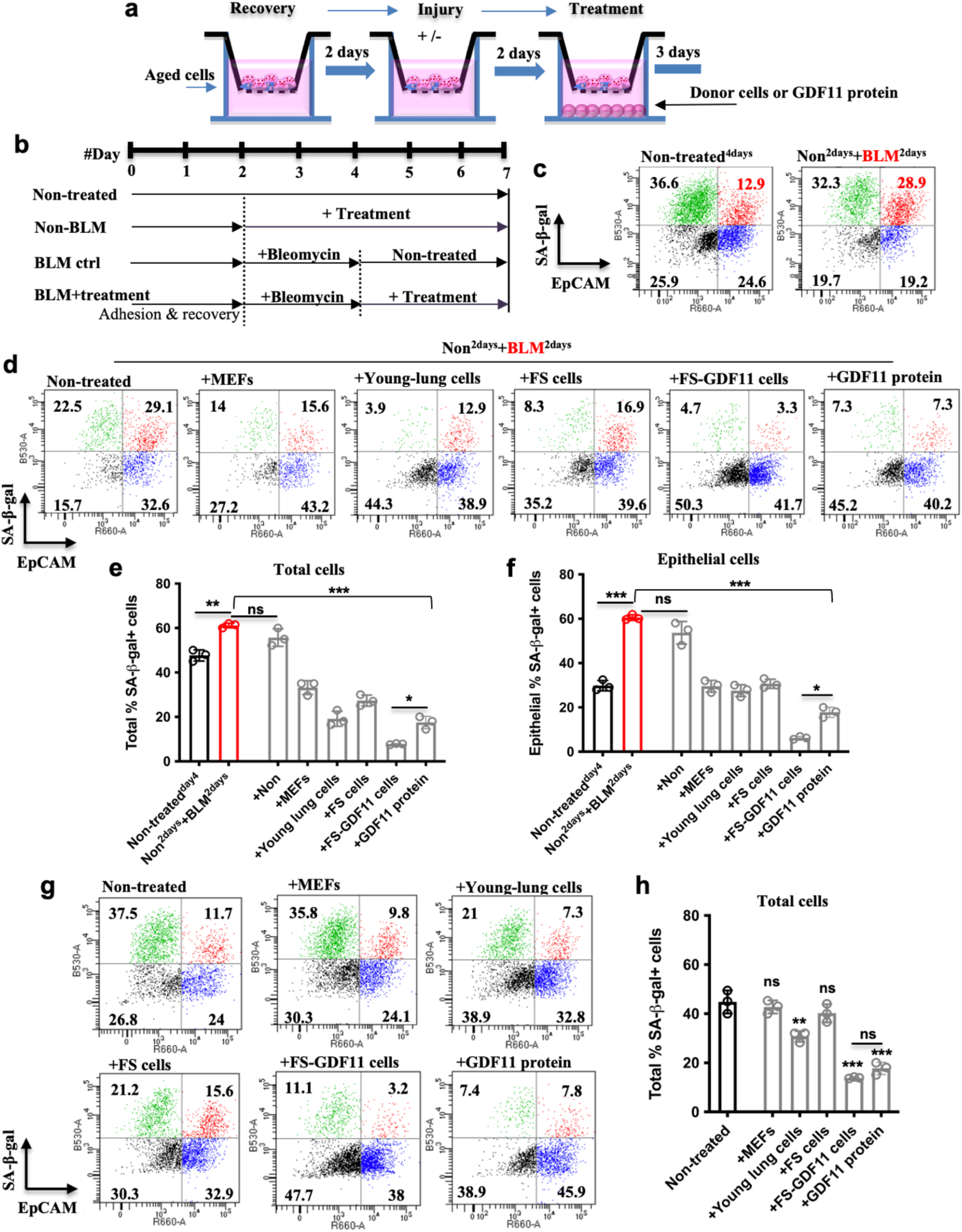
Exogenous GDF11 alleviates age-related and BLM-induced aggravated cellular senescence. **a** and **b,** Experimental scheme of the *in-vitro* study to evaluate the effect of therapeutic cell secreted GDF11 on age-related and BLM-induced aggravated cellular senescence. **c,** Representative flow cytometry dot plots of SA-β-gal and EpCAM expression levels showing BLM-induced aggravated cellular senescence in distal epithelial cells. **d,** The effect of donor cells and rGDF11 on the senescent (SA-β-gal) population. Quantification of senescence SA-β-gal+ cells in total **e** and epithelial **f** populations obtained from different groups. Representative flow cytometry dot plots **g** and quantification **h** showing the effect of donor cells and rGDF11 on the senescence of non-injured cells. *p < 0.05; **p < 0.001; ***p < 0.0001. In **c**, **d** and **g**, data are representative of a minimum of three independent biological replicates.

Previous studies found^39,41^ that BLM-induced severe cellular senescence resulted in a significant increase in SA-β-gal expression in cells, particularly in the alveolar epithelial compartment marked by EpCAM expression (Fig. 4c, e, f). Compared to control, treatments with rGDF11 or FS-GDF11^+dox^ donor cells after BLM injury demonstrated anti-senescence effects, where the latter even surpassed the rGDF11 effect. On the other hand, the treatment with FS cells, MEFs, and young lung cells showed only a moderate reduction in SA-β-gal activity (Fig. 4d, e, f). No significant changes were observed in SA-β-gal activity in cells treated with FS or MEF cells compared to the non-treated controls. Treatments with young lung cells were moderately effective compared to the rGDF11 protein and FS-GDF11^+Dox^ cell-treated groups. The remarkable senescence reversal of these two treatments with GDF11 was equally efficient (Fig. 4g, h).

### 5. Transplanted lung progenitor cells with the regulatable secretion of GDF11 accelerate fibrosis resolution in aged mice

In this proof-of-principle study, we used the BLM model in aged mice to investigate the effect of transplanted GDF11-expressing, FS-GDF11 cell-derived, Nkx2.1+ lung progenitors on fibrosis resolution. For controls, Nkx2.1+ lung progenitors were derived from the FS line, which was marked with constitutively expressed GFP, while in the FS-GDF11 line, the GFP was linked to the doxycycline-inducible GDF11 expression (Fig. 5a).

**Figure 5.**
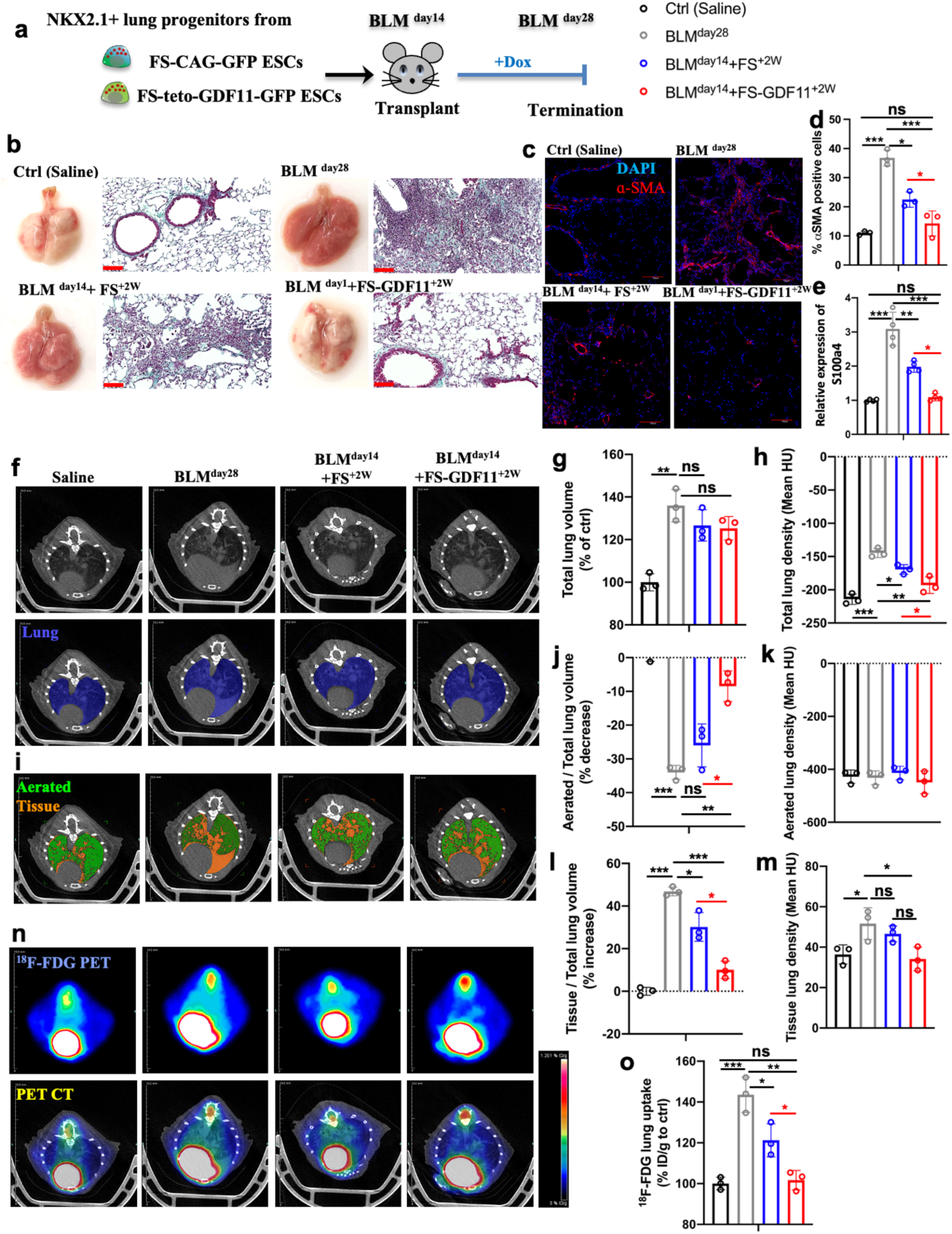
*In vivo* validation of the therapeutic utility of engineered cells to fibrosis **a** Experimental scheme of the *in vivo* study to evaluate the effect of therapeutic cell secreted GDF11 on fibrosis resolution in the mouse IPF model; **b** Representative images of whole lung appearance (left panel) and Masson’s trichrome staining (right panel) from all the experimental groups after 28 days of BLM-induced pulmonary fibrosis; **c** Representative confocal microscopy images of distal airways of all the experimental lungs showing nuclear stain DAPI (blue) and αSMA (red); **d** Quantification of αSMA positive cells; **e** The expression of S100a4 in the lungs of different groups 28 days after saline or BLM treatment, as measured by qRT-PCR comparing fold differences in gene expression in non-injured healthy controls; **f** Representative micro-CT scan images and segmentation of the lung component of different groups; Quantifications of the total lung volume **g** and density **h**; **i** Segmentation of aerated (green) and tissue (orange) components within the lungs; Quantifications of changes in aerated and tissue lung volume (**j** and **l**) and lung density (**h** and **k**); **n** Representative PET-CT images showing a remarkable improvement of fibrosis resolution in the lungs treated with engineered cells; **o** Quantification of 18F-FDG uptake in all experimental groups showing a significant reduction of 18F-FDG uptake in the lungs treated with engineered cells. *p < 0.05; **p < 0.001; ***p < 0.0001. Scale bar, 100 μm (**c**).

The mouse model of BLM-induced lung injury can be divided into three phases. The first is the inflammatory phase, which typically peaks around post-BLM day 7, followed by the fibrotic phase starting around post-BLM day 14 after the first phase subsides. The third phase, fibrosis resolution, shows a significant age-dependent delay in onset and progression^52^.

First, we investigated whether doxycycline, which has known anti-inflammatory effects, affected fibrosis resolution when administered at the onset of the fibrotic phase (post-BLM day 14). The fibrosis resolution in mice remained unaffected, as assessed at day 28 post-BLM-induced injury (Fig. S3), consistent with a previous study^67^.

At the onset of the fibrotic phase (BLM^day14^), we introduced Nkx2.1+ progenitor cells to injured aged mice transtracheally. Prior to transplantation, the Nkx2.1+ progenitors were treated with GCV *in-vitro* as a safety measure to eliminate highly proliferative potentially tumorigenic cells by the FS system and then purified based on their surface expression of carboxypeptidase M (CPM)^68^.

Animals treated solely with saline or BLM alone, without cell transplantation, served as the negative and positive controls, respectively. The animals were continuously monitored *in vivo* and sacrificed at post-BLM day 28 for further assessments (Fig. 5a). The lungs of mice treated with FS-GDF11-derived cells and kept on a dox diet for two weeks after (designated as BLM^day14^+FS-GDF11^+2W^) showed significant improvement in their external appearance compared to that of the FS-ESC-derived donor cell treated lungs (BLM^day14^+FS^+2W^). The results from *in vivo* imaging validation were consistent with the findings from histological analysis, which revealed that the GDF11-expressing cell-treated lungs had normal alveolar architecture, fewer fibrotic lesions, and less collagen deposition in the parenchyma (Fig. 5b, S4a). Immunostaining analysis of myofibroblast marker αSMA further indicated that the fibrosis was efficiently resolved in GDF11 expression cell-treated lungs (Fig. 5c, d). Accordingly, the expression of S100a4, a marker correlated with fibrosis and its progression, was also significantly reduced (Fig. 5e).

Then, micro-CT scans were utilized to conduct qualitative and quantitative assessments of fibrosis. To do this, we segmented the lung (shown as a blue overlay) within the thoracic cavity to measure total lung volume and mean density (Fig. 5f). BLM-induced inflammation and progressive fibrosis resulted in an increase in total lung volume compared to saline controls, which remained even after cell treatment (Fig. 5g). However, lungs treated with both groups of donor cells showed decreased mean lung density compared to sham-treated BLM lungs. Notably, a significant improvement was observed in the lungs treated with GDF11-expressing cells. The density value was similar to those observed in the saline-control lungs (Fig. 5h).

As demonstrated previously, improved lung function does not always accompany normalization of the enlarged total lung volume^69^. To gain further insights, the lungs were further segmented into aerated (shown as green overlay) and tissue lung contents (shown as orange overlay) using thresholding operations, and their respective proportions were quantified for comparison (Fig. 5i). The analysis revealed bulk changes in air and tissue contents of the lungs as a whole following the cell treatments. Treatment with GDF11-expressing cells yielded more prominent results, showing a significant increase in aerated lung volume (Fig. 5j), a decrease in tissue lung volume (Fig. 5l) and tissue lung density (Fig. 5m), compared to the modest beneficial effect of non-GDF11 transgenic cells in alleviating fibrosis.

^18^F-FDG PET-CT imaging has become increasingly popular in recent times for the detection of lung fibrosis and addressing the effectiveness of anti-fibrotic therapies^70^. The increased uptake of ^18^F-FDG in the lung is due to the upregulation of glucose transporters on myofibroblasts during the fibrotic phase. Our findings were consistent with previous publications, showing a significant increase in ^18^F-FDG uptake in the BLM-injured lungs compared to healthy controls. However, after cell treatments, we observed a reduction in ^18^F-FDG uptake in the injured lungs, with a greater reduction observed in the lungs treated with GDF11-expressing donor cells (Fig. 5n, o).

### 6. GDF11-expressing donor cells attenuate senescence in aged lungs, achieving fibrosis resolution

Persistent fibrosis is a condition where there is a significant buildup of fibroblasts that are senescent and resistant to apoptosis. Eliminating these senescent fibroblasts can potentially improve lung function and overall physical health in aged mice with BLM-induced lung injury^41^. Additionally, in both human IPF and BLM-injured aged mice^39–41,71^, age-related senescence in AEC-II cells has been linked to impaired fibrosis resolution in the lungs. Therefore, it is crucial to target and eliminate senescent features in both fibroblasts and AEC-II cells to develop effective therapies against pulmonary fibrosis, potentially halting or reversing its progression in aged populations.

Our *in vitro* study has shown that GDF11-expressing donor cells have a positive impact on age-related and BLM-induced cellular senescence in fibroblasts and AEC-II cells (Fig. 3d-f). Here, we investigated whether these anti-senescence beneficial effects also exist long-term in *in vivo* applications.

First, we examined donor cell retention after transplantation. To quantify retention, genomic DNA for GFP was measured using a standard curve. The results showed that the retention rates of both donor cell types were similar, 19.4±2.7% and 21.5±4.0%, respectively (Fig. 6a).

**Figure 6.**
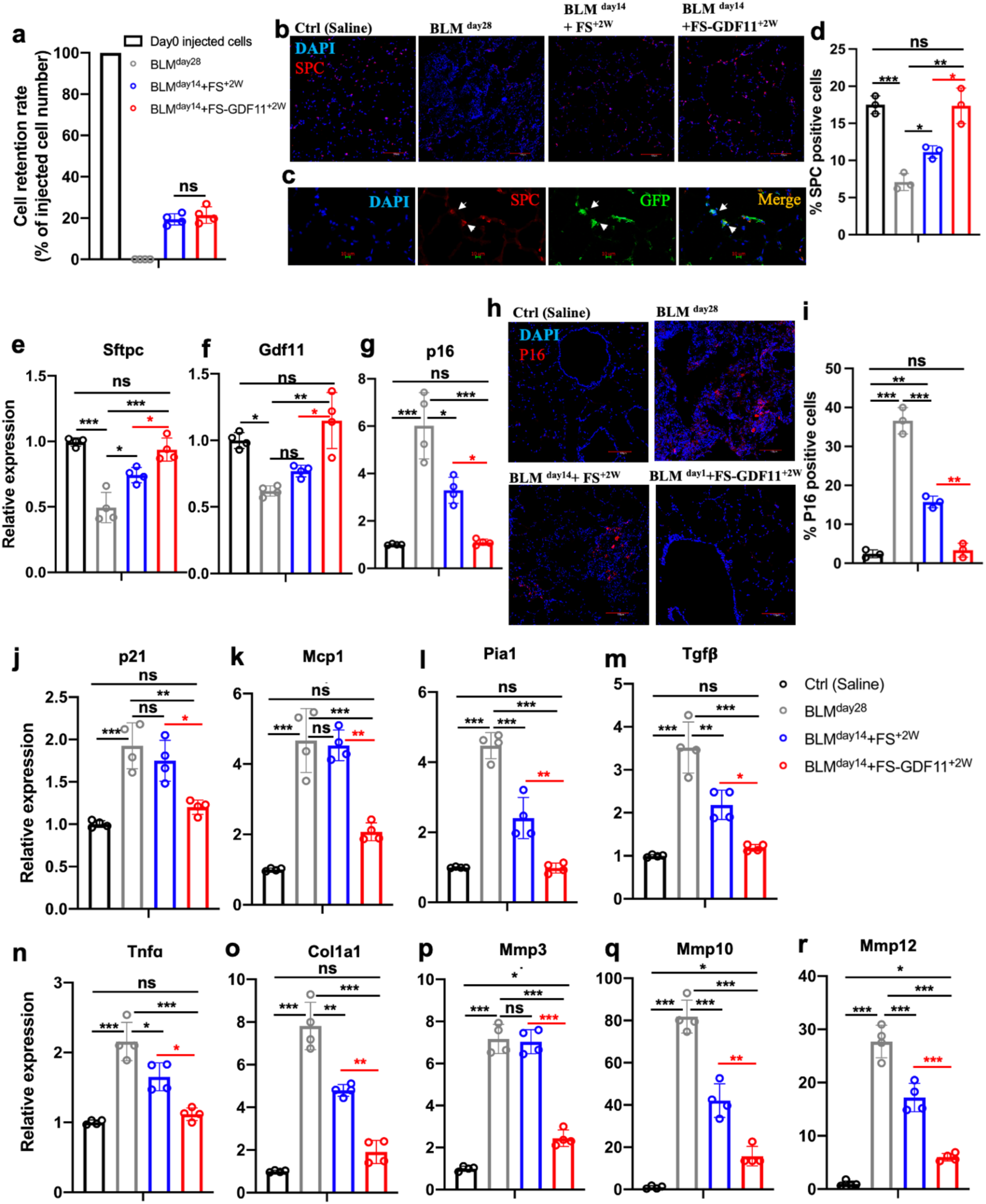
*In situ* restoration of GDF11 levels by cell transplant attenuates age-associated senescence and leads to successful fibrosis resolution **a** Retention rate of the delivered GFP+ cells in the recipient lungs (percentage of day 0 injected cells) was calculated using genomic GFP measured at day 28 of BLM treatment (relative to β-actin GFP lungs) measured by PCR; **b** Representative confocal microscopy images of distal airways of all the experimental lungs showing nuclear stain DAPI (blue) and SPC (red); **c** Representative images of the alveolar epithelium of GDF11-GFP cell treated BLM injured animals showing nuclear stain DAPI (blue), GFP (green), and SPC (red); **d** Quantification of αSMA positive cells; The expression of Sftpc **e**, Gdf11 **f** and p16 **g** in the lungs of different groups 28 days after saline or BLM treatment, as measured by qRT-PCR comparing fold differences in gene expression in non-injured healthy controls; **h** Confocal microscopy images of distal airways of all the experimental lungs showing nuclear stain DAPI (blue) and p16 (red) and **i** quantification of p16-positive cells; Transcript levels of P21 **j** and proinflammatory and profibrotic SASP factors **k**-**r**, including Mcp1, Pai1, Tgfß, Tnfα, Col1a1, Mmp3, Mmp10 and Mmp12 as measured by qRT-PCR comparing fold differences in the expression in non-injured healthy controls. *p < 0.05; **p < 0.001; ***p < 0.0001. Scale bar, 100 μm (**b** and **h**), 10 μm (**c**).

After BLM treatment, there were fewer AEC-II cells (as marked by SPC) in comparison to the untreated lungs. However, the lungs that were treated with donor cells showed an elevated number of SPC+ cells, with an especially abundant presence in the GDF11-expressing donor cell-treated group (Fig. 6b, d). The presence of GFP-expressing donor SPC+ cells demonstrated their direct contribution to the AEC-II lineage (Fig. 6c). This was further supported by the efficient restoration of the SPC encoding mRNA (Sftpc) levels (Fig. 6e).

Only the lungs treated with GDF11-expressing cells showed restoration of GDF11 mRNA levels, which were comparable to those seen in healthy controls (Fig. 6f). This indicates that the GDF11-expressing cells were effective in providing GDF11 *in situ*.

We observed that the elevated transcriptional activation of p16 induced by BLM was downregulated in the lungs treated with donor cells. This decrease was more pronounced with GDF11-expressing donor cells, reaching levels comparable to that of healthy controls (Fig. 6g). Immunostaining also revealed that the number of p16+ senescent cells was significantly reduced in these lungs (Fig. 6h, i).

Similar to p16, the cell cycle inhibitor p21 is also implicated in cellular senescence^72^. We observed a significant elevation in p21 expression in the lungs of BLM-injured mice compared to the control group. However, significant downregulation of p21 was only evident in the lungs treated with GDF11-expressing cells (Fig. 6j). This finding indicates that the restoration of GDF11 activity can also eliminate p21-dependent cell senescence, which may facilitate alveolar regeneration^73^.

Furthermore, we conducted a study on the senescence-associated secretory phenotype (SASP) components in the lungs of all experimental groups. The level of transcripts of proinflammatory and profibrotic SASP factors, including Mcp1, Pai1, Tgfß, Tnfα, Col1a1, Mmp3, Mmp10, and Mmp12, were measured by RT-PCR (Fig. 6k-r). Both groups of donor cells showed positive effects. Despite lacking exogenous GDF11, the control donor cells were able to reduce the expression of most of these genes except for Mcp1 and Mmp3 (Fig. 6k, p). Remarkably, the GDF11-expressing cells showed a greater potential in downregulating all these genes, including Mcp1 and Mmp3. Additionally, we observed no obvious adverse effects in the mice that received exogenous GDF11 from transplanted GDF11-expressing cells (Fig. S4b).

## Discussion

We developed a combination of cell and gene therapy approaches to target human IPF using a mouse model of the disease. For the gene therapy component, we chose GDF11 inducible expression because of the implication of its regenerative potential in certain physiological functions. For the cell therapy component, we generated lung progenitors from engineered ESCs with controlled proliferation by implementing our FS system^49^ into the cells. Upon delivery to injured aging lungs, these progenitor cells can engraft and restore the damaged alveolar epithelium. These cells also serve as a “factory on-site” and can produce the necessary amount of GDF11 to compensate for the natural decline of this factor resulting from aging or lung-related diseases. This approach effectively mitigates age-associated senescence, which is further exacerbated in diseased conditions, resulting in the successful resolution of age-related persistent fibrosis.

Some of the controversy surrounding the function of GDF11 is rooted in its high level of homology with myostatin (GDF8)^32,74^. Both GDF11 and myostatin are members of the TGF-β superfamily and are involved in tissue regeneration, playing critical roles in regulating cellular processes. They have been extensively studied for their potential therapeutic applications in regenerative medicine. However, their specific regenerative utilities differ significantly. While myostatin primarily regulates muscle growth and restricts muscle stem cell proliferation^32,75^, GDF11 demonstrates a broader regenerative capacity beyond the muscle. GDF11 has been shown to promote neurogenesis in the brain, improve cardiovascular function, and facilitate tissue repair across various organs, such as the liver and kidney^76–83^. On the other hand, myostatin’s effects are primarily focused on skeletal muscle development and maintenance^84–86^. Therefore, GDF11’s diverse multi-tissue regenerative properties make it a more appealing candidate for comprehensive therapies, particularly in the context of lung health. Our research indicates a link between decreased levels of GDF11 and impaired fibrosis resolution associated with cellular senescence in aging. Through both *in vitro* and *in vivo* studies, we have demonstrated the regenerative potential of GDF11 in the lung, suggesting broader implications for diseases mediated by age-related senescence^87^, such as IPF.

The role of GDF11 in fibrosis is a subject of ongoing research and remains somewhat contentious due to its membership in the BMP subfamily of the TGF-β superfamily. The TGF-β subfamily members, such as β1, β2, and β3, are known to be strongly associated with fibrosis by promoting the accumulation of extracellular matrix components and contributing to tissue scarring^88^. While GDF11 is generally thought to have anti-fibrotic effects by promoting BMP signaling^48,89,90^, it is important to note that its role in fibrosis can be complex and context-dependent. It can activate both the SMAD1/5/8 pathway, which counterbalances TGF-β signaling in epithelial-mesenchymal transition (EMT) and fibrosis, and the profibrotic SMAD2/3 pathway^19^. Some studies have suggested that GDF11 may also have profibrotic effects in certain conditions or tissues^91–93^. The dual roles of GDF11 in fibrosis may be influenced by factors such as the cellular environment, the activity of other signaling pathways, and the specific disease context.

Hence, an appropriate dosage is essential for harnessing the anti-fibrotic potential of GDF11 to specific tissue and disease phenotypes. However, this process could be costly and arduous when using recombinant protein, as demonstrated by numerous studies involving intraperitoneal injection of rGDF11^44,94–96^. High-dose or long-term exposure to rGDF11 can impede therapeutic outcomes and lead to severe adverse effects, such as neurotoxicity, cachexia and mortality^44,46,128^. These limitations favour using transplanted cells as a localized source of GDF11 production, targeting specific organs or sites of regeneration as needed.

To address the challenges associated with the limited practicality of rGDF11, we have developed a cell-based treatment strategy that allows for localized and sustained release of GDF11. This approach involves utilizing transplanted lung progenitor cells that have been differentiated from FailSafe ES cells further engineered to express GDF11 in a controlled manner. We transplanted these via transtracheal delivery rather than systemic intravenous injection. Animals treated with the cell therapy showed a complete recovery in body weight. Additionally, whole-body PET-CT scans showed no signs of tumours or adverse effects (Fig S4b).

In the current study, we chose GDF11 as a prototype regenerative factor. While the existing viewpoint suggests that GDF11 may not be the sole universal facilitator of regeneration or rejuvenation, our methodology provides a flexible approach that can be adapted for various other regenerative and anti-aging agents, such as sirtuins^1–6^, Klotho^7–13^, insulin-like growth factor 1 (IGF-1)^14–17^, and platelet factor 4 (PF4)^18^. For all of these agents, our approach has the potential to address practical limitations associated with their use, such as high cost, short half-life, and the need for precise dosing to effectively address specific aging and disease-related characteristics.

Despite the importance of this proof-of-concept study, certain areas warrant further investigation to enhance therapeutic effectiveness. While senescence was notably reduced by GDF11-expressing cell treatment, the suppression of matrix metalloproteinases (MMPs) genes (including Mmp3, Mmp10, and Mmp12) associated with extracellular matrix (ECM) degradation and fibrosis resolution was not fully achieved and these genes remained elevated compared to healthy controls. Therefore, additional research is necessary to optimize *in vivo* GDF11 expression, including the dosing of transplanted cells, dox feed and timing/duration of induction, in order to improve therapeutic efficacy.

We demonstrate the potential of GDF11-expressing lung progenitor cells in cell-based therapy for ameliorating age-related IPF in a mouse model. The capacity to deliver GDF11 *in situ* through regulated transgene expression enables a more precise and targeted strategy to address cellular senescence and fibrosis. The effectiveness of this method could have wider implications for treating fibrotic lung conditions linked to aging and senescence.

There is an increasing number of clinical trials exploring cell therapy for treatments of degenerative diseases. These trials use cells that are either derived from autologous or from allogeneic pluripotent stem cells^97^. Regarding targeting lung diseases, protocols have been developed to generate human lung resident cells with directed *in vitro* differentiation from induced pluripotent stem cells^98–102^. However, to advance toward readiness for stem cell-based therapies, two major challenges need to be addressed: ensuring cell therapy safety and promoting acceptance of allogeneic cells without requiring immune suppression of patients. We have proposed genome editing solutions for these challenges and developed the FailSafe cell system^49^ and the induced allogeneic cell tolerance (iACT)^103^ systems. The integration of these two systems could lead to the creation of safe and universally applicable off-the-shelf therapeutic products that are accessible to all humans. This advancement could significantly expand the scope of diseases that cell therapy can effectively target, offering a new treatment avenue for degenerative diseases on a larger scale within the field of medicine.

## Methods

### Animal husbandry

C57BL/6N (Stock No 005304) mice were purchased from Jackson Laboratory. 6-8 week-old and 12-14 month-old mice were used for BLM injury and cell replacement therapy studies. Animals were maintained as an in-house breeding colony under specific pathogen-free conditions. All animal care protocols and procedures were performed in accordance with relevant guidelines and with approval by the Institutional Animal Care and Use Committee of the University Health Network (Toronto, Ontario, Canada).

### Bleomycin administration and cell delivery

For BLM-induced injury, BLM 2.5 U kg^-1^ body weight was administered intratracheally. Donor cells (10^6^ cells in 50μl PBS) were delivered intratracheally 14 days after injury. Control animals received the same volume of saline without any cells. The mice receiving cells were rotated to ensure equal dispersion of the cell suspension to both lungs.

### Construction of piggyBac vectors

The plasmid containing the cDNA of murine Gdf11 was obtained from the Lunenfeld-Tanenbaum Research Institute Open Freezer repository. To prepare the constructs for expression, the cDNAs were inserted into piggyBac transposon expression vectors using the Gateway cloning kit (Thermo Fisher 12535029) according to the manufacturer’s protocol. In summary, the Gdf11 gene was amplified by extension PCR using PrimeStar HS master mix (Takara R040), with gateway-compatible attb1/attb2 sites added to the 5’ and 3’ ends, respectively. The resulting product was then recombined into the gateway pDONr221 vector (Thermo Fisher 12536017), and the transgene insertion was confirmed by sequencing the resulting entry vector using flanking M13 Forward and M13 Reverse primers. Subsequently, the transgene-containing entry clone was introduced into piggyBac destination vectors via Gateway cloning to produce the final vectors for transgene expression in cells. At each stage of cloning, vectors were introduced into chemically competent DH5alpha cells (Invitrogen), followed by the selection of colonies on LB agar plates supplemented with either kanamycin (Sigma K1377) or ampicillin (Sigma 10835242001). Colonies were then cultured in LB broth (Wisent 809-060-L), and bacterial plasmid DNA was extracted using the Presto Mini Bacterial DNA kit (Geneaid PDH300).

### Fluorescence-activated cell sorting and analysis

For extracellular staining, freshly isolated or cultured cells were suspended and incubated in 0.5% (vol/vol) FBS-PBS with an optimally pre-titered mixture of antibodies and relevant isotype controls for approximately 30 minutes on ice. The labelled cells were then washed and re-suspended at a concentration of 3∼5 × 10^6^ cells/mL in 0.5% (vol/vol) FBS-PBS. Cell viability was assessed by propidium iodide or 4,6-diamidino-2-phenylindole (DAPI; Sigma) staining at a concentration of 1μg/mL. For intracellular antigen analysis, cells were fixed and stained using a Fix and Perm kit (Invitrogen) following the manufacturer’s instructions. Sorting was conducted using a MoFlo BRU cell sorter (Becton Dickinson), acquisition was performed using a BD LSRII analyzer (Becton Dickinson), and data were analyzed using FlowJo software.

### Immunofluorescence

Samples were fixed with 4% paraformaldehyde (PFA) for 30 minutes and subsequently blocked with a solution containing 5% goat serum and 2% BSA in PBS supplemented with 0.5% Triton X-100 for 1 hour. Primary antibodies, diluted in BSA/PBS, were then applied to the samples and left to incubate overnight at 4°C. Following this, secondary antibodies, such as AlexaFluors 488, 532, 546, 633, or 647 (Invitrogen), were applied based on the species for which the primary antibody was raised and incubated for 2 hours at room temperature. Nuclear staining was achieved using 2 mg/mL of 4,6-diamidino-2-phenylindole (DAPI; Sigma). Stained samples were mounted with immunofluorescent mounting medium (DAKO), and appropriate non-specific IgG isotypes were utilized as controls. Immunoreactivities of antigens were visualized as single optical planes using an Olympus Fluoview confocal microscope and analyzed with FV10-ASW 2.0 Viewer software.

### Real-time PCR analysis

Total RNA was extracted from cells or lung tissue utilizing the RNeasy Kit (Qiagen) following the manufacturer’s guidelines. Subsequently, cDNA synthesis and analysis were conducted using Superscript III (Sigma) in accordance with the manufacturer’s instructions. Differential gene expression was assessed using SYBR green detection (Roche). Real-time PCR reactions were performed in triplicate for each sample. Housekeeping genes were used to standardize gene expression levels. Normalized mRNA levels were presented relative to the corresponding controls.

### PET/CT Imaging

#### Mouse preparation and dose administration for imaging

Mice were subjected to overnight fasting while having unrestricted access to water. An intravenous injection of a net dose ranging from 9-12 MBq of fluorine-18 fluorodeoxyglucose (^18^F-FDG) was administered via the tail vein catheter, followed by a saline flush. Subsequently, mice were anesthetized with 1.5-2% isoflurane and 1-1.5L/min oxygen. PET imaging was initiated approximately 1-hour post-injection.

#### Imaging

PET/CT imaging was performed on a Mediso NanoScan® SPECT/CT/PET system (Budapest, Hungary). The CT scan parameters included 50 kVp, 980 µA, and a 300 ms exposure time, followed by a 10-minute PET acquisition. Mediso NanoScan Nucline software (Version 3.04.012.0000) was used for image acquisition and reconstruction (including automatic PET and CT co-registration). CT scans were reconstructed using the medium voxel and thin slice thickness settings, resulting in an isotropic voxel size of 78 µm x 78 µm x 78 µm. PET images were reconstructed using the standard normal Tera-Tomo 3D reconstruction setting (OSEM-based for 4 subsets, 4 iterations, with attenuation and scatter correction, isotropic voxel size of 400 µm).

#### Image Analysis

Image processing, as well as manual and semi-automatic segmentation of the lungs and heart, was performed using Siemens Inveon Research Workplace (IRW) 4.2 (Siemens Medical Solutions, Tennessee, USA).

### Statistics

Statistical analysis was performed using GraphPad Prism 5.0 statistical software (San Diego, CA, USA). The statistical significance of multiple groups was compared to each other using Tukey’s multiple comparison test ANOVA. A p-value of <0.05 was considered significant.

## Supplemental Information

Supplemental information includes 4 figures and 2 tables.

## Acknowledgements

This study was supported by the Canadian Institutes of Health Research Foundation Grant [143231], Canada Research Chair [950-230422] to AN, and Grosman-Lubin Fellowship Award to LG. We would like to thank the Spatio-Temporal Targeting and Amplification of Radiation Response (STTARR) program and its affiliated funding agencies, with special thanks to Deborah Scollard, Maria Bisa (from the UHN Animal Resources Centre), Dr. Luke Kwon and Teesha Komal for imaging support.

## Competing Interests

A.N. is a shareholder and co-founder of panCELLa Inc. The company obtained an exclusive license for the commercialisation of the FailSafe^TM^ technology. The other authors declare no competing financial interests and non-financial interests.

## Author Contributions

L.G. designed experiments, performed experiments, analyzed data and wrote the manuscript. P.D. assisted with *in vivo* animal injury models and cell delivery. E.S. performed cell injection of teratoma assay. E.D.J. assisted with cell culture. C.L. monitored teratoma formation. T.K.W contributed to the *in vivo* study design and edited the manuscript. A.N. generated the hypothesis, designed experiments, provided funding and wrote the manuscript.

## Notes

### Competing Interest Statement

The authors have declared no competing interest.

