## Supplementary material for "GDF11 secreting cell transplant efficiently ameliorates age-related pulmonary fibrosis": Supplemetary Data

### Supplementary figures

#### Figure S1

**a**

| Gene symbol | Gene name | Cellular function |
|---|---|---|
| Gapdh | glyceraldehyde-3- phosphate dehydrogenase | Catalyzes the reversible oxidative phosphorylation of glyceraldehyde-3-phosphate in glycolysis |
| B2m | β-2-microglobulin | Beta-chain of major histocompatibility complex class I molecules. Involved in immune response. |
| Eef2 | Eukaryotic Translation Elongation Factor 2 | Protein Synthesis |
| Hprt | Hypoxanthine phosphoribosyltransferase 1 | Purine synthesis through the purine salvage pathway |
| Rpl13a | Ribosomal protein L13a | Structural component of the large 60S ribosomal subunit |
| Ppia | Peptidylprolyl isomerase A | Protein coding, a cyclosporin binding-protein |

**b**

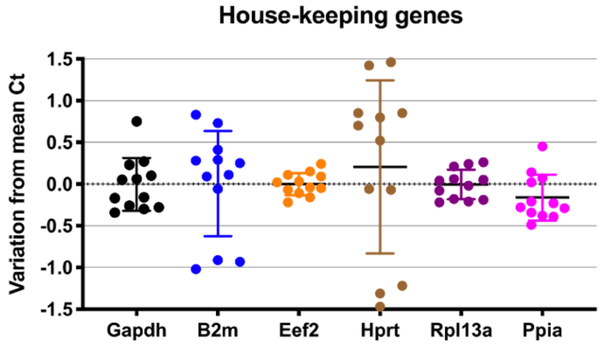


**c**

| | Summary | P Value |
|---|---|---|
| **Gapdh** | | |
| Young (Saline) vs. Young (Bleomycin) | ns | 0.2606 |
| Young (Saline) vs. Old (Saline) | ns | 0.2195 |
| Young (Saline) vs. Old (Bleomycin) | ns | 0.1833 |
| Young (Bleomycin) vs. Old (Saline) | ns | 0.9997 |
| Young (Bleomycin) vs. Old (Bleomycin) | ** | 0.0016 |
| Old (Saline) vs. Old (Bleomycin) | ** | 0.0012 |
| **B2m** | | |
| Young (Saline) vs. Young (Bleomycin) | **** | <0.0001 |
| Young (Saline) vs. Old (Saline) | ns | 0.9685 |
| Young (Saline) vs. Old (Bleomycin) | ns | 0.1295 |
| Young (Bleomycin) vs. Old (Saline) | **** | <0.0001 |
| Young (Bleomycin) vs. Old (Bleomycin) | **** | <0.0001 |
| Old (Saline) vs. Old (Bleomycin) | ns | 0.2968 |
| **Eef2** | | |
| Young (Saline) vs. Young (Bleomycin) | *ns* | *0.5409* |
| Young (Saline) vs. Old (Saline) | ns | 0.2694 |
| Young (Saline) vs. Old (Bleomycin) | ns | 0.9213 |
| Young (Bleomycin) vs. Old (Saline) | ns | 0.9601 |
| Young (Bleomycin) vs. Old (Bleomycin) | ns | 0.8923 |
| Old (Saline) vs. Old (Bleomycin) | ns | 0.6271 |

| | | |
|---|---|---|
| **Hprt** | | |
| Young (Saline) vs. Young (Bleomycin) | **** | <0.0001 |
| Young (Saline) vs. Old (Saline) | * | 0.0102 |
| Young (Saline) vs. Old (Bleomycin) | * | 0.039 |
| Young (Bleomycin) vs. Old (Saline) | **** | <0.0001 |
| Young (Bleomycin) vs. Old (Bleomycin) | **** | <0.0001 |
| Old (Saline) vs. Old (Bleomycin) | **** | <0.0001 |
| **Rpl13a** | | |
| Young (Saline) vs. Young (Bleomycin) | ns | 0.7793 |
| Young (Saline) vs. Old (Saline) | ns | 0.9994 |
| Young (Saline) vs. Old (Bleomycin) | ns | 0.1458 |
| Young (Bleomycin) vs. Old (Saline) | ns | 0.7115 |
| Young (Bleomycin) vs. Old (Bleomycin) | ns | 0.6148 |
| Old (Saline) vs. Old (Bleomycin) | ns | 0.1146 |
| **Ppia** | | |
| Young (Saline) vs. Young (Bleomycin) | ns | 0.8497 |
| Young (Saline) vs. Old (Saline) | ns | 0.4109 |
| Young (Saline) vs. Old (Bleomycin) | ns | 0.6997 |
| Young (Bleomycin) vs. Old (Saline) | ns | 0.8761 |
| Young (Bleomycin) vs. Old (Bleomycin) | ns | 0.2436 |
| Old (Saline) vs. Old (Bleomycin) | ns | 0.0519 |

**Figure S1.** Selection of house-keeping genes that are stably expressed in the lungs during aging in both physiological and pathological conditions **a** The list of candidate house-keeping genes; **b** The plots represent variation from the mean RT-qPCR threshold (Ct) values of candidate housekeeping genes starting from equal amounts of RNA obtained from lung tissues of young and old mice at day 28 post saline and BLM-administration, as measured by qRT-PCR; **c** The summary of statistic analysis.

**Figure S2**

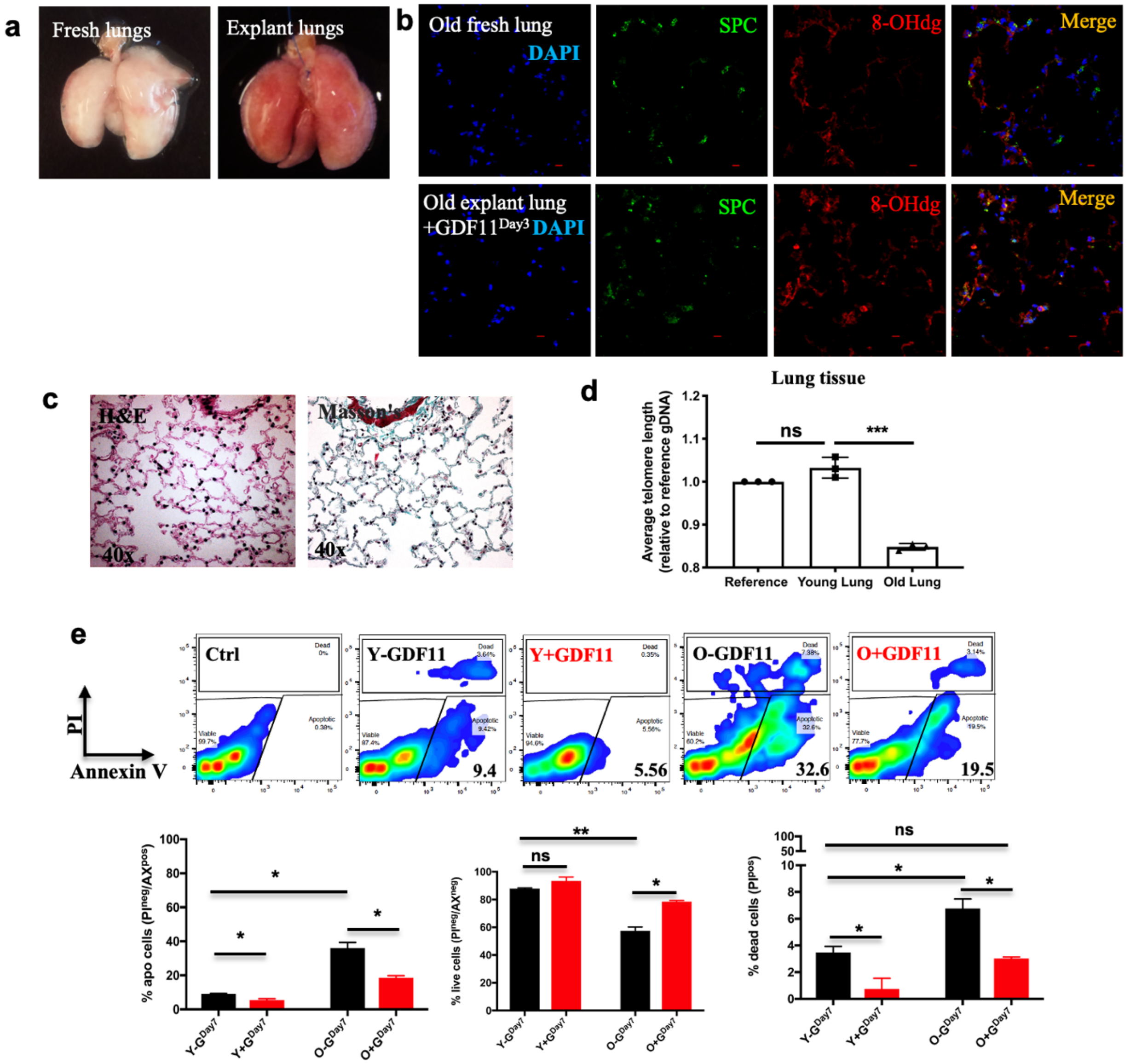


**Figure S2.** Exogenous GDF11 can partially ameliorate age-related cellular deterioration in the distal lung **a** Representative images of fresh and explant lungs; **b** Representative confocal microscopy images of distal airways of explant lungs of old mice treated with or without GDF11 recombinant protein for 3 days showing nuclear stain DAPI (blue), SPC (green) and 8-OHdG (red); **c** Representative images of Hematoxylin and Eosin and Masson's trichrome staining of the explant lungs treated with GDF11 recombinant protein for 7 days; **d** Expression levels of average telomere length in lung tissues obtained from young and old mice each, as measured by qRT-PCR comparing fold differences in the expression in reference tissues of 8-12 weeks mice; **e** Flow cytometric analysis of PI and Annexin V expression in cells cultured in each condition, and quantification of live, dead and apoptotic cells. *p < 0.05; **p < 0.001; ***p < 0.0001. In **d**, data are representative of a minimum of three independent biological replicates. Scale bar, 10 μm (**b**).

**Figure S3**

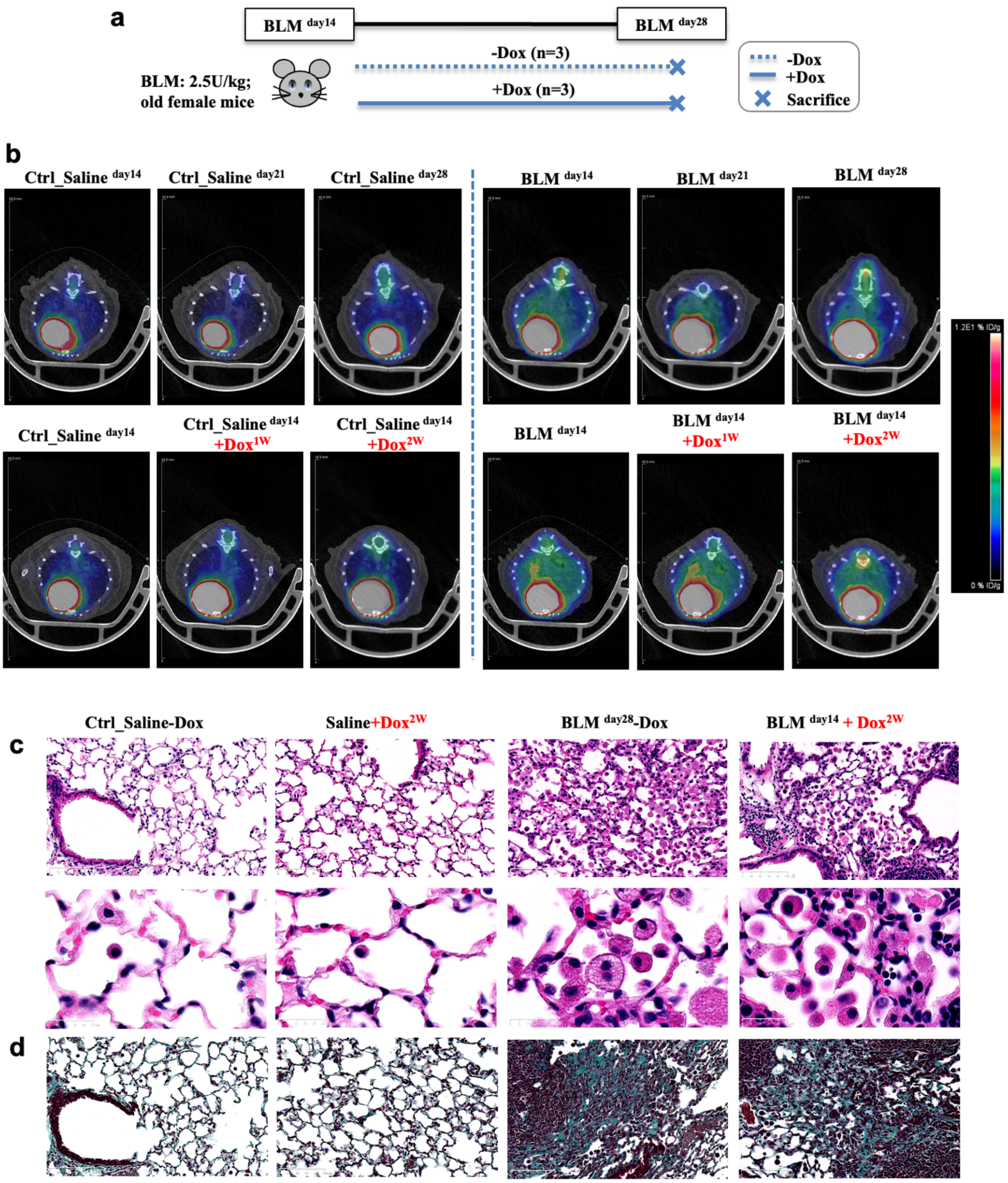

**Figure S3.** Established fibrosis in aged mice was not affected by the doxycycline diet employed for in-*vivo* transgene activation **a** The experimental scheme of the *in-vivo* study to evaluate the effect of doxycycline diet on fibrosis resolution in the mouse IPF model; **b** Representative PET-CT scan images showing $^{18}$F-FDG uptake in the lungs of different groups; Hematoxylin–eosin **c** and Masson's trichrome **d** stained of lung sections showed inflammatory cell infiltration and collagen deposition in BLM-induced fibrotic lungs, regardless of regular or doxycycline diet.

Figure S4

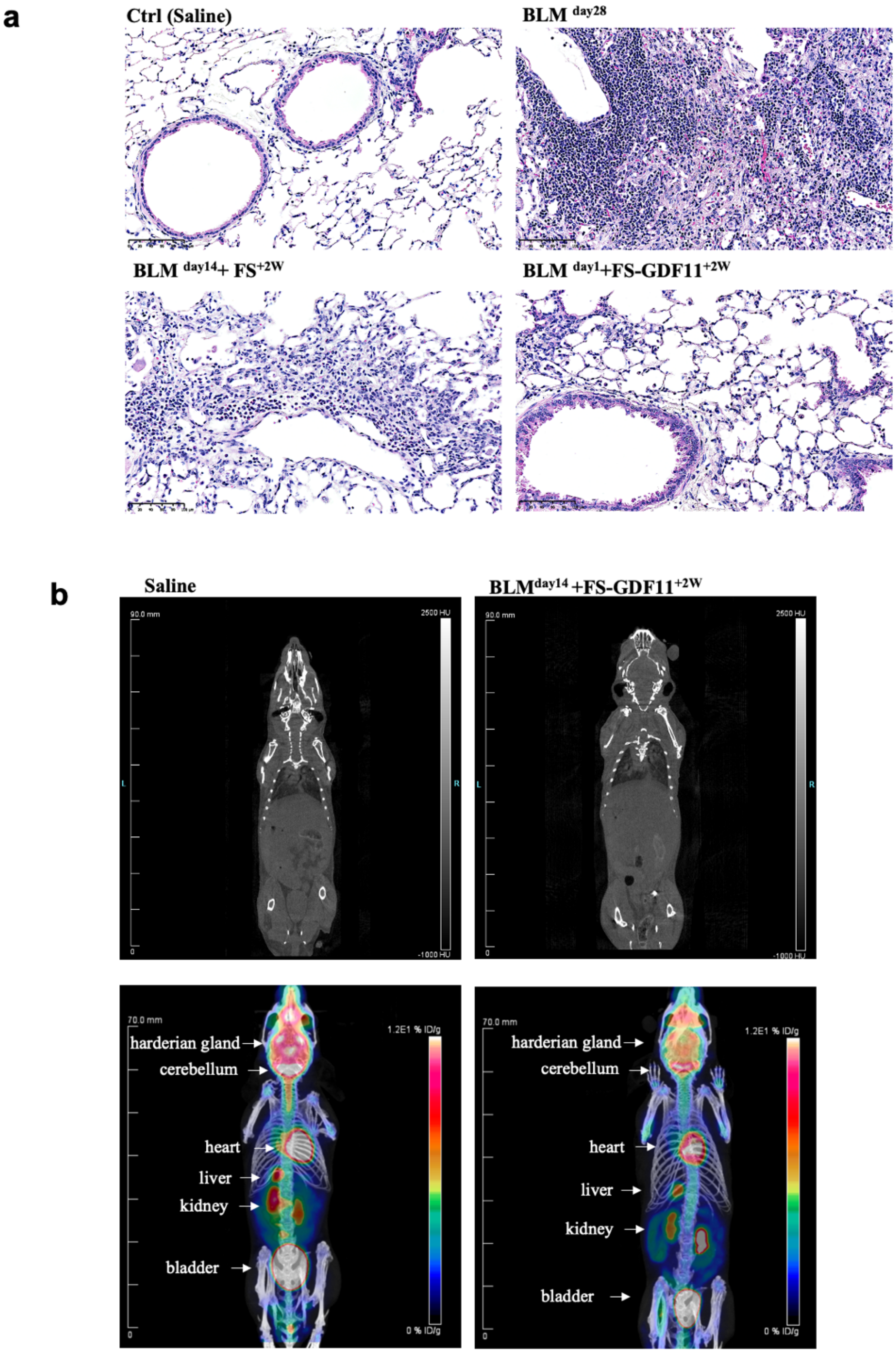


**Figure S4.** No obvious adverse effects were observed in the mice that received exogenous GDF11 from transplanted designer cells **a** Representative images of Hematoxylin–eosin staining from all the experimental groups after 28 days of BLM-induced pulmonary fibrosis; **b** Representative whole-body micro-CT (top panel) and PET-CT (bottom panel) scan images of non-injured healthy control mice and GDF11 cell-treated injured recipients ($BLM^{day14}$ +$FS\text{-}GDF11^{+2W}$ ) showing an expected biodistribution of radiotracer with activity in normal tissue and physiologic radiotracer excretion. Scale bar, 100 µm (**a**).

**Table 1.** List of antibodies used in the studies

| Antibody name | Company | Cat. # | Validation |
|---|---|---|---|
| c-Kit | Thermofisher | 47-1172-82 | ES and ES-derived cells |
| CXCR4 | Biolegend | 146508 | ES and ES-derived cells |
| GDF11 | Cedarlane | orb1183719 | Adult lung tissue; |
| GDF11 | Cedarlane | orb101175 | ES and ES-derived cells |
| 8-OHdg | Abcam | ab62623 | Adult lung tissue |
| $\gamma$H2AX | Abcam | ab11174 | Adult lung tissue and cells |
| CD45 | BD Pharmingen | 553081 | Adult lung cells |
| CD31 | BD Pharmingen | 553373 | Adult lung cells |
| EpCAM | Abcam | ab95641 | Adult lung cells; ES-derived cells |
| Nanog | Thermofisher | 50-5761-82 | ES cells |
| | Abcam | ab80892 | ES cells |
| Oct4 | Abcam | ab184665 | ES cells |
| NKX2.1 | Abcam | ab242428 | ES-derived cells |
| SPC | Millipore | AB3786 | Adult lung tissue |
| GFP | Thermo Fisher | A-21311 | Adult lung tissue |
| P16 | Abcam | ab54210 | Adult lung tissue and cells |
| a-SMA | Abcam | ab124964 | Adult lung tissue |

**Table 2.** Tabulated list of all qPCR primers used in the studies

| Gene name | Primer sequence 5'-3' | | Length |
|---|---|---|---|
| Gdf11 | Forward | CCGGCGTCACATCCGTATC | 19 |
| | Reverse | ACTTGCTTGAAGTCGATGCTC | 21 |
| S100a4 | Forward | TCCACAAATACTCAGGCAAAGAG | 23 |
| | Reverse | GCAGCTCCCTGGTCAGTAG | 19 |
| p16 | Forward | CTCTGCTCTTGGGATTGGC | 19 |
| | Reverse | GTGCGATATTTGCGTTCCG | 19 |
| Gapdh | Forward | AGGTCGGTGTGAACGGATTTG | 20 |
| | Reverse | TGTAGACCATGTAGTTGAGGTCA | 21 |
| B2m | Forward | TGACCGGCTTGTATGCTATC | 20 |
| | Reverse | CAGTGTGAGCCAGGATATAG | 20 |
| Eef2 | Forward | TGTCAGTCATCGCCCATGTG | 19 |
| | Reverse | CATCCTTGCGAGTGTCAGTGA | 20 |
| Hprt | Forward | AGCAGGTCAGCAAAGAACT | 19 |
| | Reverse | CCTCATGGACTGATTATGGACA | 22 |
| Rpl13a | Forward | CTCAAGGTCGTGCGTCTGAA | 20 |
| | Reverse | TGGCTGTCACTGCCTGGTACT | 21 |
| Ppia | Forward | GGGTTCCTCCTTTCACAGAA | 20 |
| | Reverse | GATGCCAGGACCTGTATGCT | 20 |
| p21 | Forward | CGGTGTCAGAGTCTAGGGGA | 18 |
| | Reverse | ATCACCAGGATTGGACATGG | 24 |
| Gadd45b | Forward | CGGCCAAACTGATGAATGT | 21 |
| | Reverse | TCTGCAGAGCGATATCATCC | 23 |
| Atf3 | Forward | CTCTGGCCGTTCTCTGGA | 24 |
| | Reverse | GGTCGCACTGACTTCTGAGG | 22 |
| Il6 | Forward | TCCTTAGCCACTCCTTCTGT | 20 |
| | Reverse | AGCCAGAGTCCTTCAGAGA | 19 |
| Mmp13 | Forward | GGACTCACTGTTGGTCCCTG | 20 |
| | Reverse | GGATTCCCGCAAGAGTCACA | 20 |
| Sftpc | Forward | GCAAAGAGGTCCTGATGGAG | 20 |
| | Reverse | GCAGTAGGTTCCTGGAGCTG | 20 |
| Mcp1 | Forward | AACTACAGCTTCTTTGGGACA | 21 |
| | Reverse | CATCCACGTGTTGGCTCA | 18 |
| Pia1 | Forward | CGTGTCAGCTCGTCTACAG | 19 |
| | Reverse | CTATGGTGAAACAGGTGGACT | 21 |
| Tgf**β** | Forward | CCGAATGTCTGACGTATTGAAGA | 23 |
| | Reverse | GCGGACTACTATGCTAAAGAGG | 22 |
| Tnfα | Forward | TCTTTGAGATCCATGCCGTTG | 21 |
| | Reverse | AGACCCTCACACTCAGATCA | 20 |
| Col1a1 | Forward | CATTGTGTATGCAGCTGACTTC | 22 |
| | Reverse | CGCAAAGAGTCTACATGTCTAGG | 23 |
| Mmp3 | Forward | TGTGGAGGACTTGTAGACTGG | 21 |
| | Reverse | GATGAACGATGGACAGAGGATG | 22 |
| Mmp10 | Forward | TGTTGCTCTTCAGTATGTGTGT | 22 |
| | Reverse | CCAGGAATTGAGCCACAAGT | 20 |
| Mmp12 | Forward | GCTCCTGCCTCACATCATAC | 20 |
| | Reverse | GGCTTCTCTGCATCTGTGAA | 20 |
